# Dynamic parallel transmit diffusion MRI at 7T

**DOI:** 10.1101/2024.01.06.574463

**Authors:** Minghao Zhang, Belinda Ding, Iulius Dragonu, Patrick Liebig, Christopher T. Rodgers

## Abstract

Diffusion MRI (dMRI) is inherently limited by SNR. Scanning at 7T increases intrinsic SNR but 7T MRI scans suffer from regions of signal dropout, especially in the temporal lobes and cerebellum. We applied dynamic parallel transmit (pTx) to allow whole-brain 7T dMRI and compared with circularly polarized (CP) pulses in 6 subjects.

Subject-specific 2-spoke dynamic pTx pulses were designed offline for 8 slabs covering the brain. We used vendor-provided B_0_ and B_1_^+^ mapping. Spokes positions were set using the Fourier difference approach, and RF coefficients optimized with a Jacobi-matrix high-flip-angle optimizer. Diffusion data were analyzed with FSL.

Comparing whole-brain averages for pTx against CP scans: mean flip angle error improved by 15% for excitation (2-spoke-VERSE 15.7° *vs* CP 18.4°, *P*=0.012) and improved by 14% for refocusing (2-spoke-VERSE 39.7° *vs* CP 46.2°, *P*=0.008). Computed spin-echo signal standard deviation improved by 14% (2-spoke-VERSE 0.185 vs 0.214 CP, *P*=0.025). Temporal SNR increased by 5.4% (2-spoke-VERSE 8.47 *vs* CP 8.04, P=0.004) especially in the inferior temporal lobes. Diffusion fitting uncertainty decreased by 6.2% for first fibres (2-spoke VERSE 0.0655 vs CP 0.0703, *P*<0.001) and 1.3% for second fibers (2-spoke VERSE 0.139 *vs* CP 0.141, *P*=0.01). In conclusion, dynamic parallel transmit improves the uniformity of 7T diffusion-weighted imaging. In future, less restrictive SAR limits for parallel transmit scans are expected to allow further improvements.

## Introduction

Diffusion MRI (dMRI) provides quantitative information about the Brownian motion of water molecules within tissue through application of diffusion encoding gradients.[1, 2] The mean diffusivity and the fractional anisotropy[3] are important biomarkers for tissue microstructure.[4] In the context of neuroimaging, diffusion MRI has been applied to the evaluation of ischemic stroke[5] and tumors[6], to study human brain connectivity[7] and to understand the mechanisms of neurodegenerative disorders such as Alzheimer’s disease and Parkinson’s disease.[8–10]

dMRI is intrinsically limited by low signal-to-noise ratios (SNR) since the diffusion weighting stems from signal loss due to the diffusion gradients. As a result, in a clinical setting, diffusion MRI scans are usually acquired with lower resolution than structural scans.[11] Although scanning at higher field strength improves the intrinsic SNR per unit scan time[12], the accompanying decrease in tissue T_2_ poses challenges for the acquisition protocols.[13] Furthermore, scanning at ultra-high field introduces well-known challenges of increased transmit field (B_1_^+^) inhomogeneity and increased RF power deposition (i.e. specific absorption rate, SAR).

Previous studies have shown that imaging at 7T makes it feasible to acquire high resolution (1.05 mm isotropic) dMRI images[14], and that *static* parallel transmission (pTx) – also known as “RF shimming” – reduces transmit field inhomogeneities and improves SNR, especially in the lower parts of the brain.[15]

Static pTx allows the RF amplitude and phase of each channel to vary, whereas *dynamic* pTx allows each channel to play out a different arbitrary waveform. Dynamic pTx has been successfully implemented in many neuroimaging sequences, including structural scans[16, 17] and functional MRI BOLD scans[18] yielding improved data quality. We hypothesize that the extra degrees of freedom from dynamic pTx will also improve the quality of 7T dMRI.

In this study, we aimed to improve 7T dMRI coverage using per-subject optimized 2-spoke dynamic pTx pulses, which we compare against conventional circularly polarized (CP or “TrueForm”) pulses for whole brain diffusion tensor imaging (DTI) in volunteers.

## Material and Methods

### Equipment

Scans were performed in a 7T MAGNETOM Terra MRI scanner (Siemens Healthcare, Erlangen, Germany) running VE12U SP01 software with an 8Tx/32Rx head coil (Nova Medical, MA, USA) at the University of Cambridge. Some scans were also performed in a 7T MAGNETOM Terra MRI scanner at the University of Glasgow with a custom 8Tx/32Rx head coil.[19]

### Pulse sequence

The vendor’s single-shot echo planar imaging (EPI) sequence (‘ep2d_diff’, shown in Figure 1b) was modified to allow the choice of conventional circularly polarized (CP or “TrueForm”) pulses or 2-spoke dynamic parallel transmit pulses for excitation and refocusing.

**Figure 1.**
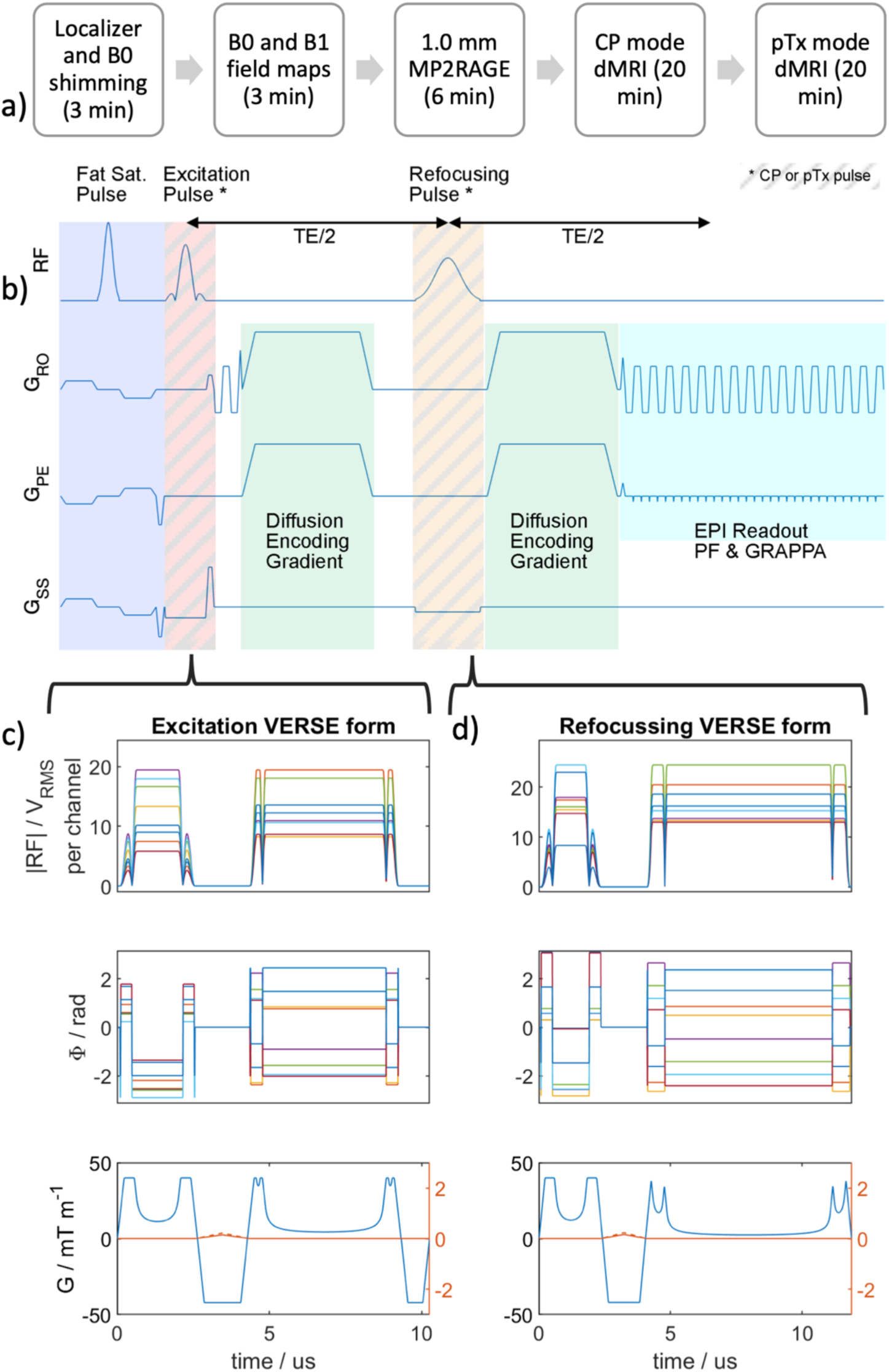
a) The pTx diffusion acquisition protocol. The pulse design was carried out during the MP2RAGE and CP mode diffusion scans and did not add extra time. b) The SS-EPI diffusion sequence diagram. CP or pTx pulses can be used for both excitation and refocusing pulses. The minimum TE is limited by the diffusion gradient and EPI train after the refocusing pulse. c) A typical pair of excitation and refocusing pulses. The pulses are converted to VERSE form to reduce SAR.

We based our reference CP protocol on the Human Connectome Project’s (HCP) published 7T dMRI protocol,[14] since the HCP consortium are acknowledged leaders in dMRI. The HCP project used MAGNETOM 7T MRI scanners (Siemens) running a previous VB17 version of software for their study, so we matched the parameters as best as we could onto the newer VE12U SP01 software of our 7T Terra scanner.

### Pulse design

Subject-specific 2-spoke pulses for pTx acquisitions were calculated offline in MATLAB (MathWorks, USA) with a modified version of the vendor’s pTx pulse design (PPD) framework. Maps of ΔB_0_ and per-channel B_1_^+^ were acquired using the vendor-provided dual-echo gradient echo (GRE) and magnetization-prepared turbo-FLASH[20] (TFL) sequences. A whole brain region of interest (ROI) mask was computed by thresholding the magnitude image from the B_0_ mapping sequence. These field maps and details of the reference voltage, prescribed slice positions and SAR information were exported in three MATLAB “.mat” files and used for offline pulse design.

We ran an offline pulse design for monopolar 2-spoke pulses for excitation and for refocusing in each slab (a total of 8 slabs covering the brain, see below). The target excitation and refocusing flip angles were 80° and 160° to reduce the specific absorption rate (SAR). A linear correction[21] was applied to the TFL B_1_^+^ maps. The last spokes were all fixed at the center of transmit k-space (“DC spokes”). The first spokes were positioned in transmit k-space (*k*_T_-space) according to the Fourier transform-based method[22] based on the field maps for the central slice in each slab. Briefly, the DC spoke complex coefficients were optimized; the resulting predicted transverse magnetization was computed using the ΔB_0_ and per-channel B_1_^+^ maps; this was subtracted from the target magnetization to give a residual; the residual was Fourier transformed to *k*_T_-space; the first spoke was placed at the *k*_T_-space point with highest residual amplitude. The complex coefficients of both spokes were then jointly optimized, initially using the magnitude least squares (MLS) algorithm[23] applying the small tip angle (STA) approximation, and using Tikhonov regularization on the L_2_ norm of the RF amplitude[24] to control SAR. MLS pulse design used the ΔB_0_ and per-channel B_1_^+^ maps for all slices in the slab but neglected T_2_* decay. Two solver and initialization combinations were used for each calculation, namely a least-squares with QR factorization (LSQR)[25] solver with CP mode initial values, and a Powell[26] solver with all-zero initial values. The superior MLS solution was then used as the starting point for the Jacobi-matrix large-tip-angle optimization with gradient descent included in the vendor’s PPD toolbox. Peak channel voltage constraints were enforced. SAR was controlled with a Tikhonov regularization factor λ fixed at 0.025. This value was found during in silico trials as the maximum energy penalty without significantly deteriorating flip angle uniformity. The RF spoke subpulses were Shinnar-Le-Roux (SLR)[27] optimized 90° and 180° pulses provided by the vendor, as used in the HCP diffusion protocol. The subpulse lengths were 3.84 ms for excitation and 5.12 ms for refocusing, corresponding to half of the pulse lengths used with the vendor’s “low SAR” sequence option.

To reduce SAR, the 2-spoke pulses were converted to variable-rate selective excitation (VERSE)[28] pulses with the minimum-time algorithm.[29] A binary search was performed for the maximum voltage, so that the total duration of the 2-spoke VERSE pulse was kept the same as the original SLR pulse and the phase relationship between the two spokes did not need to be re-optimized.

Bloch simulations of the predicted spin-echo signal amplitude were performed after the scans to evaluate the CP and pTx pulse performance. The excitation and refocusing pulse combinations were simulated with strong crusher gradients on both sides of the refocusing pulse. Simulations used 9x9x9 isochromats equally spaced spatially across the target voxel to simulate dephasing from the crusher gradients. The transverse magnetization at the time of the spin-echo was averaged for each voxel. Relaxation was ignored.

### In vivo Acquisition

Six healthy volunteers (3 male, 3 female, aged 25 – 42 years, mean age 33 years) gave written, informed consent, and the study was approved by the Cambridge Human Biology Research Ethics Committee [HBREC.2020.27]. The acquisition protocol is summarized in Figure 1a.

Whole brain B_0_ shimming was performed with the vendor’s GRE Brain method. Interactive shimming was performed where necessary to reduce the water FWHM linewidth to below 50 Hz. B_0_ and B_1_^+^ field maps were obtained after B_0_ shimming by running a short “dummy” pTx-enabled scan. The dual-echo GRE B_0_ adjustment scan were acquired at 4.4x4.4x4.4 mm^3^, and TFL B_1_^+^ scan at 4.0x4.0x5.0 mm^3^. The dummy scan was set up that the PPD framework interpolated these maps into 3.15 mm slices, so that each slice exactly corresponded to 3 slices in the diffusion acquisition (detailed below) and allowed even division of the slabs, without unnecessarily increasing the size of the pulse design problem.

T_1_-weighted structural images were acquired with MP2RAGE[30] at 1.0 mm isotropic resolution with the PASTEUR universal pulse sequence[16] in sagittal orientation. Other parameters were: 4300ms TR, 840 / 2370ms inversion times, 1.91ms TE, 250 Hz/Px bandwidth and 5°/6° nominal flip angles.

Diffusion MRI was performed with our single-shot echo planar imaging sequence using acquisition parameters based on the published 7T HCP protocol.[14] The whole brain was covered in 120 transverse slices with no gap, with 1.05 mm isotropic voxels, 71 ms TE, 1388 Hz/px bandwidth, 3x GRAPPA with GRE reference scans and 6/8 partial Fourier in plane acceleration, and 14300 ms TR. The excitation and refocusing flip angles were 90° and 180° for the CP scans but reduced to 80° and 160° to reduce SAR for the pTx scans. The vendor’s default 110° CP fat saturation pulse was used for all scans. The transmit voltage was set to achieve no greater than 15% overflip in the center of the brain and no greater than 235V, in line with the HCP recommendation.

Due to the SAR limitations with the pTx sequence (discussed below), only single-band acquisitions were performed. We acquired 30 diffusion directions in two shells (b=1000 and 2000 s/mm^2^) in one phase encoding direction (anterior-posterior, AP). Images with no-diffusion encoding (b=0 s/mm^2^) were acquired with the same parameters, in two phase encoding directions (AP and PA) to allow distortion corrections. The PA images were acquired with 10 repeats to measure temporal SNR (tSNR). All subjects were first scanned in the CP mode.

Per-subject pTx pulse optimization was performed for 8 equal-sized slabs covering the brain in Matlab (Mathworks Inc, Nattick, MA, USA, version 2022b) as described above. This took 6-7min on a desktop computer (Intel Core i9-10900X, 64 GB RAM). PTx-dMRI scans were then acquired. For some subjects, the SAR limits were still exceeded, in which case TR was increased until the scan would run.

### Repeat scans on a coil with full VOP supervision

Three subjects in the aforementioned CP – pTx comparison acquisitions were scanned again at the Imaging Centre of Excellence, Glasgow, UK. Ethical approval was provided by the College of Medical, Veterinary & Life Sciences (MVLS), University of Glasgow (Project No: 200210019). These scans also used a Magnetom Terra 7T scanner with VE12U SP01 software, but this time with a different 8Tx32Rx head coil for which full (non-diagonal) VOP supervision was enabled. All pulse design parameters were kept the same. The scanner reported SAR levels and achievable TR were compared with that those from the Nova head coil with 1W per channel and 8W total power limits in First Level mode. Data from these scans was not otherwise included in our group analyses.

### Data processing

DICOM images were converted to NIfTI format with dcm2nii[31]. Signal and tSNR comparisons were carried out using the raw b=0 no diffusion images without further processing. A Gaussian filter of standard deviation 1 was applied to the resultant mask for smoothing. Brain masks of the raw EPI images were obtained using ‘bet’ (FSL).

The voxel-wise tSNR percentage difference was calculated in the EPI space by

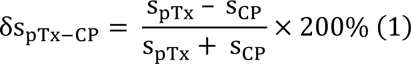

where s_pTx_ is the tSNR of the pTx image and s_CP_ is that of the CP image. To account for the TR and hence acquisition time difference in some subjects, an adjusted tSNR was calculated by dividing the pTx tSNR by 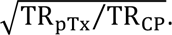

Diffusion image processing was carried out with the FSL pipeline. Susceptibility-induced off-resonance field estimation and the distortion correction of b=0 images was performed with ‘topup’.[32] Brain masks for the distortion-corrected EPI images were obtained with ‘bet’, followed by eddy current, distortion and motion correction with ‘eddy’[33]. Tensor fitting was performed with ‘dtifit’. The probabilistic diffusion model parameters were estimated with ‘bedpostx’,[34] which outputs information including the estimated mean of the fractional anisotropy, diffusivity and the orientation and its uncertainty of up to three principal fibers.

Data were registered and projected to the Montreal National Institute 152 (MNI 152) common space of 1 mm resolution via the T_1w_ MP2RAGE structural image with a linear 12 degree-of-freedom affine registration (FMRIB’s Linear Image Registration Tool, FLIRT)[35] for statistical analyses.

Statistical significance was tested with a one-tailed paired Student’s t-test with 5 degrees of freedom at a significance level of 0.05.

## Results

Figure 1c shows the RF and gradient waveforms of a pair of excitation and refocusing monopolar 2-spoke pTx pulses. We report TE based on the center of the second (DC) spoke.

Table 2 shows pulse optimization results for all subjects in every slice. The whole brain flip angle RMSE decreased significantly with pTx vs CP. For the excitation pulse with target flip angle 80°, mean RMSE improved (15.7° 2-spoke VERSE *vs* 18.4° CP; 15% reduction; *P*=0.012). For the refocusing pulse with target flip angle 160°, mean RMSE also improved (39.7° 2-spoke VERSE *vs* 46.2°; 14% reduction; *P*=0.008). Due to the increased sensitivity to ΔB_0_ from VERSE conversion, in some subjects, the slices immediately above the frontal sinus showed slice bending. The pTx scans are SAR restricted, and the SAR level and the final TR used are recorded in Table 1.

**Table 1.**
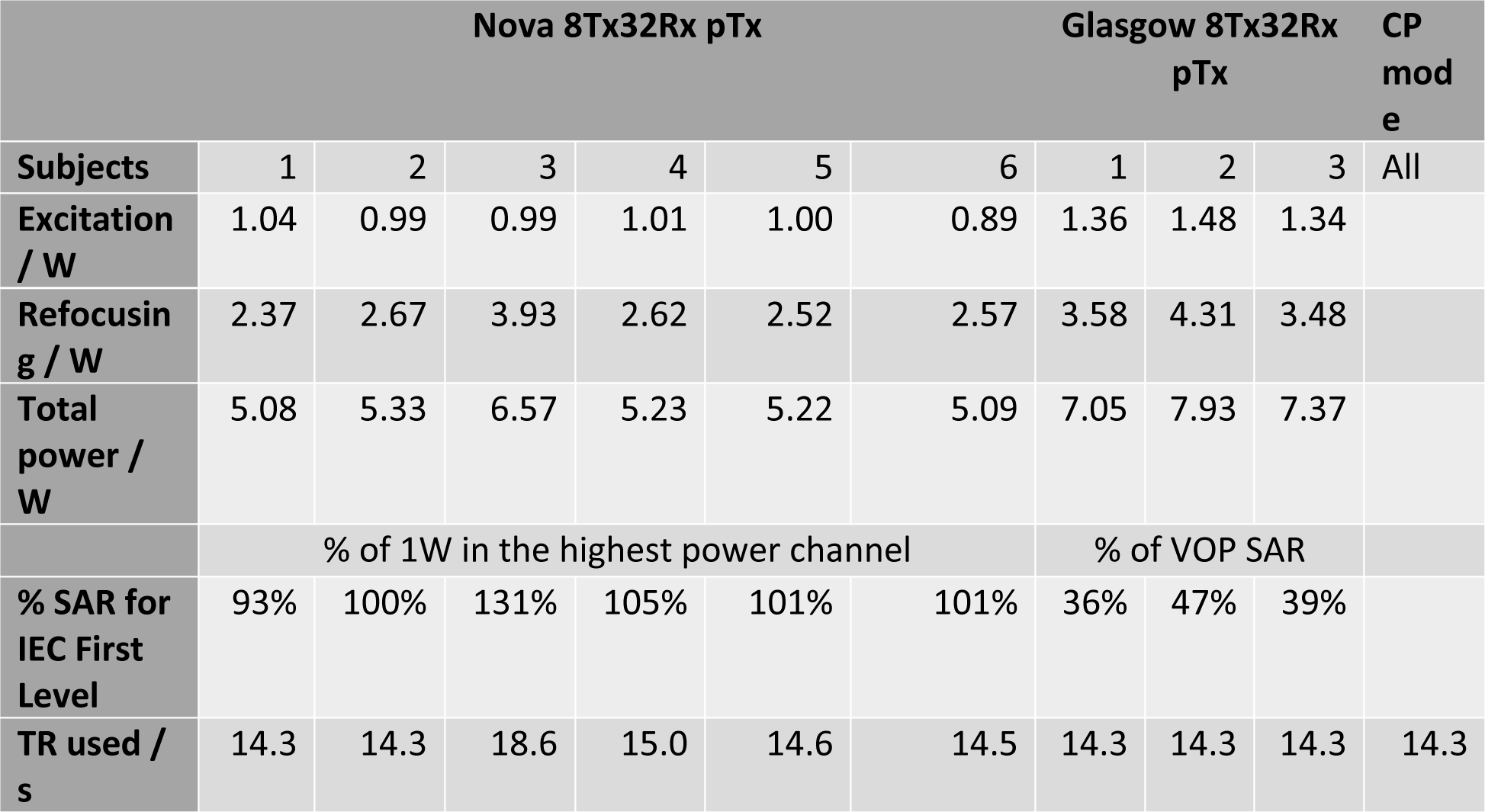
The pTx 2-spoke pulse powers in all subjects. In pTx mode on 7T Terra systems with VE12U software version, the IEC first level SAR limit for the current ‘diagonal’ VOP model for the Nova 8Tx32Rx coils is 1 W per channel, 8 W total. This is compared with the full VOP supervision with a custom 8Tx32Rx head coil at University of Glasgow. Powers are calculated for TR=14.3s for 120 slices, but where the SAR limit is exceeded, the actual TR used for the scans was increased accordingly (last row).

**Table 2.**
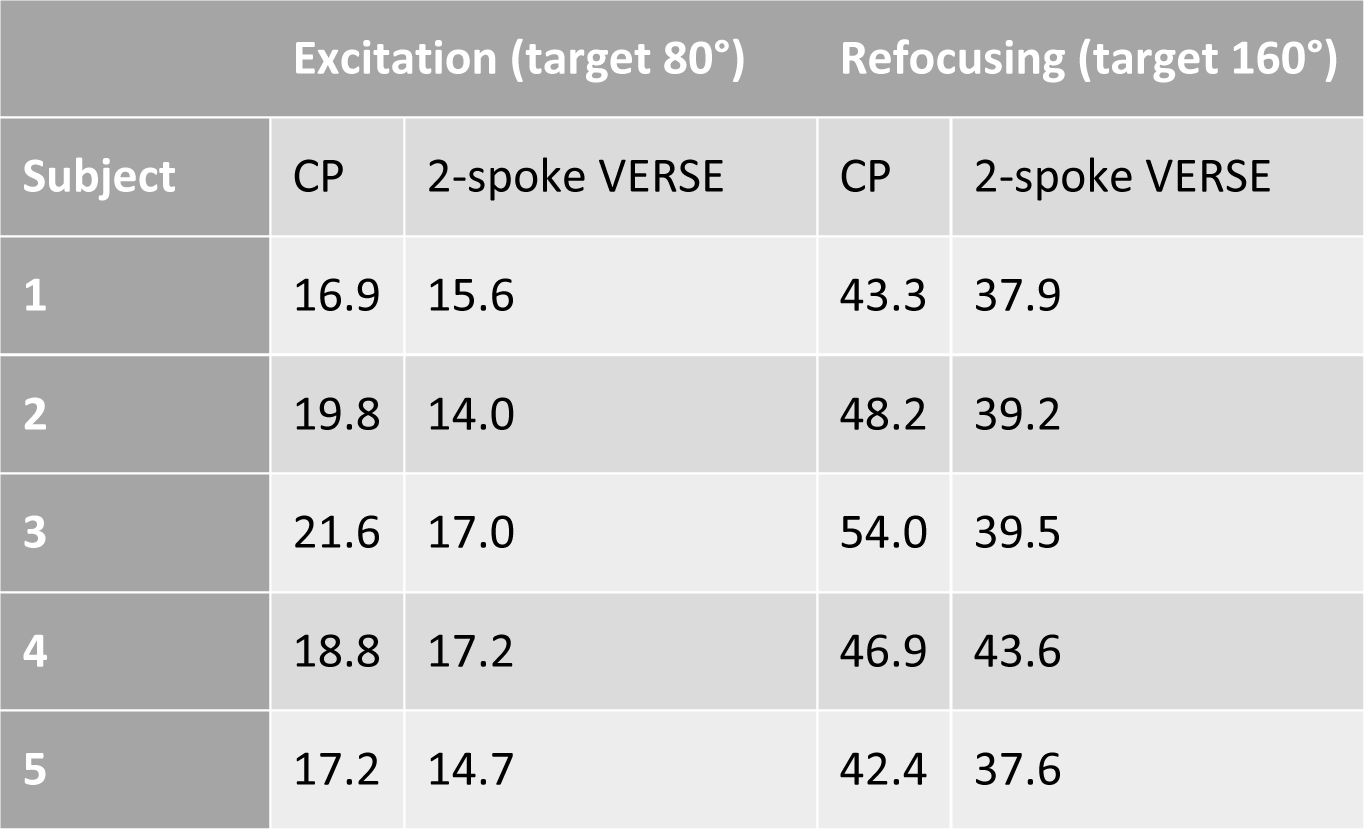
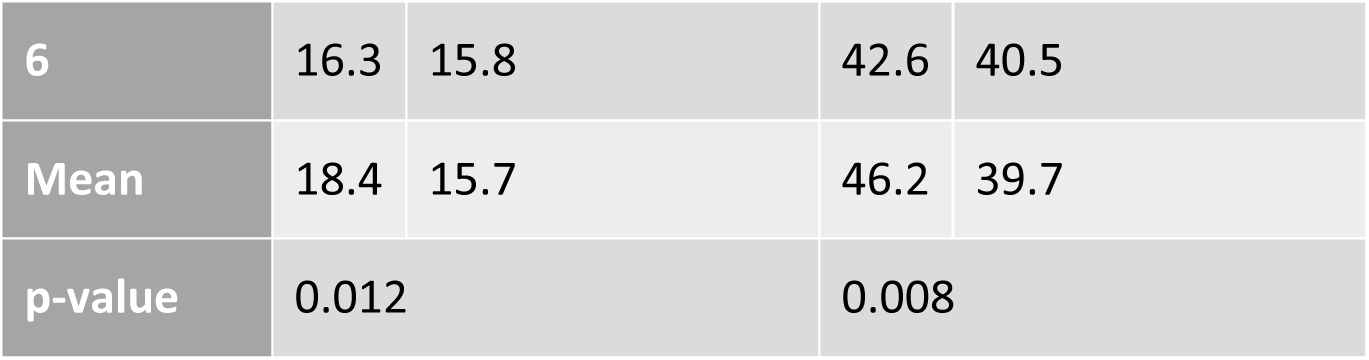
Whole brain flip angle root mean squared errors for all subjects by Bloch simulation. Errors are calculated within a brain mask retrospectively obtained from the structural image with ‘bet’. The decrease in flip angle RMSE with pTx 2-spoke VERSE for both excitation (P=0.012) and refocusing (P<0.008) pulses are significant compared to CP in paired t-tests.

Figure 2 (a-c) compares the simulated spin-echo signal intensity of the CP and 2-spoke pulses in one subject, masked with a transformed brain mask from the MP2RAGE image to highlight signal changes inside the brain. As expected at 7T, the CP pulses showed a strong central brightening effect but gave weak signals in the inferior temporal lobes, occipital lobes, and the cerebellum. The pTx 2-spoke VERSE pulses produced more uniform transverse magnetization across the whole brain, although with a slightly decreased signal intensity in the center of the brain compared to CP scans. The standard deviation of the transverse magnetization inside the brain mask decreased by 14% for the inter-subject mean (2-spoke VERSE 0.185 vs 0.214 CP, P=0.025).

**Figure 2.**
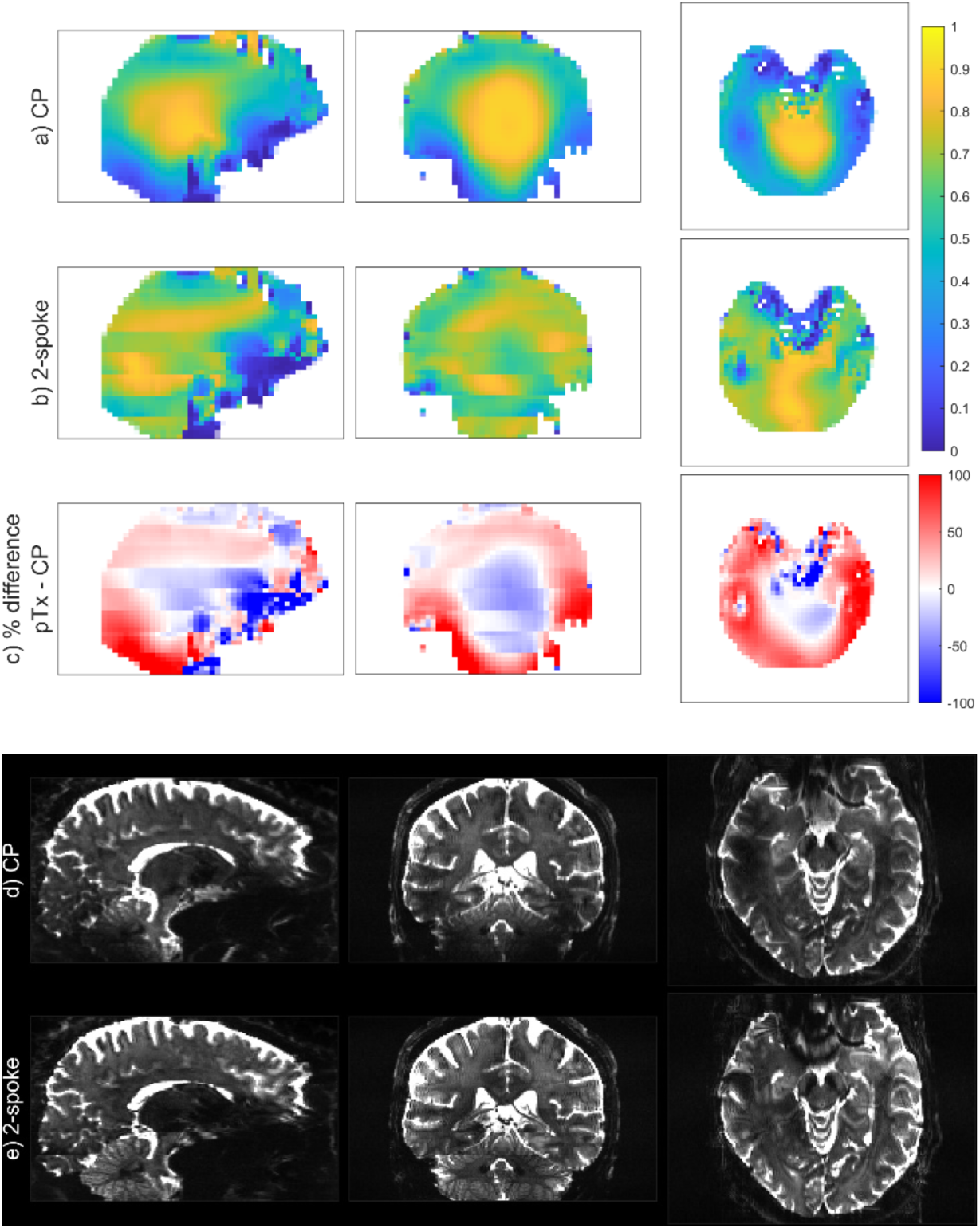
(a-c) Bloch simulaton of the spin-echo transverse magnetization comparison between CP and 2-spoke VERSE pulses in one subject. The brain mask was obtained with FSL ‘bet’ with the MP2RAGE image, to highlight the changes inside the brain and to calculate statistics. (d-e) The raw b=0 image comparison in the same subject between the circularly polarized (CP) mode acquisition and the 2-spoke pTx acquisition. The same windowing was applied to all images.

Figure 2 (d-e) compares the raw signal intensity between the CP and the 2-spoke acquisitions of the same slices. Consistent with the Bloch simulation results, the CP images shows signal dropouts in the inferior parts of the temporal lobes and in the cerebellum. The pTx 2-spoke pulses produced more signal in these areas which results in a visibly more uniform intensity profile in the coronal view. Note that the pTx images show intensity variations at the transition between slabs due to their using different sets of pulses.

The pTx pulses improved temporal SNR (tSNR) in the cerebellum (Figure 3). Over the whole brain mask, the mean tSNR of CP and 2-spoke acquisitions are 8.04 and 8.64 respectively. After correcting for the lengthened TR in some subjects, the adjusted pTx 2-spoke tSNR is 8.47, representing a 5.4% increase from the CP images (P=0.004).

**Figure 3.**
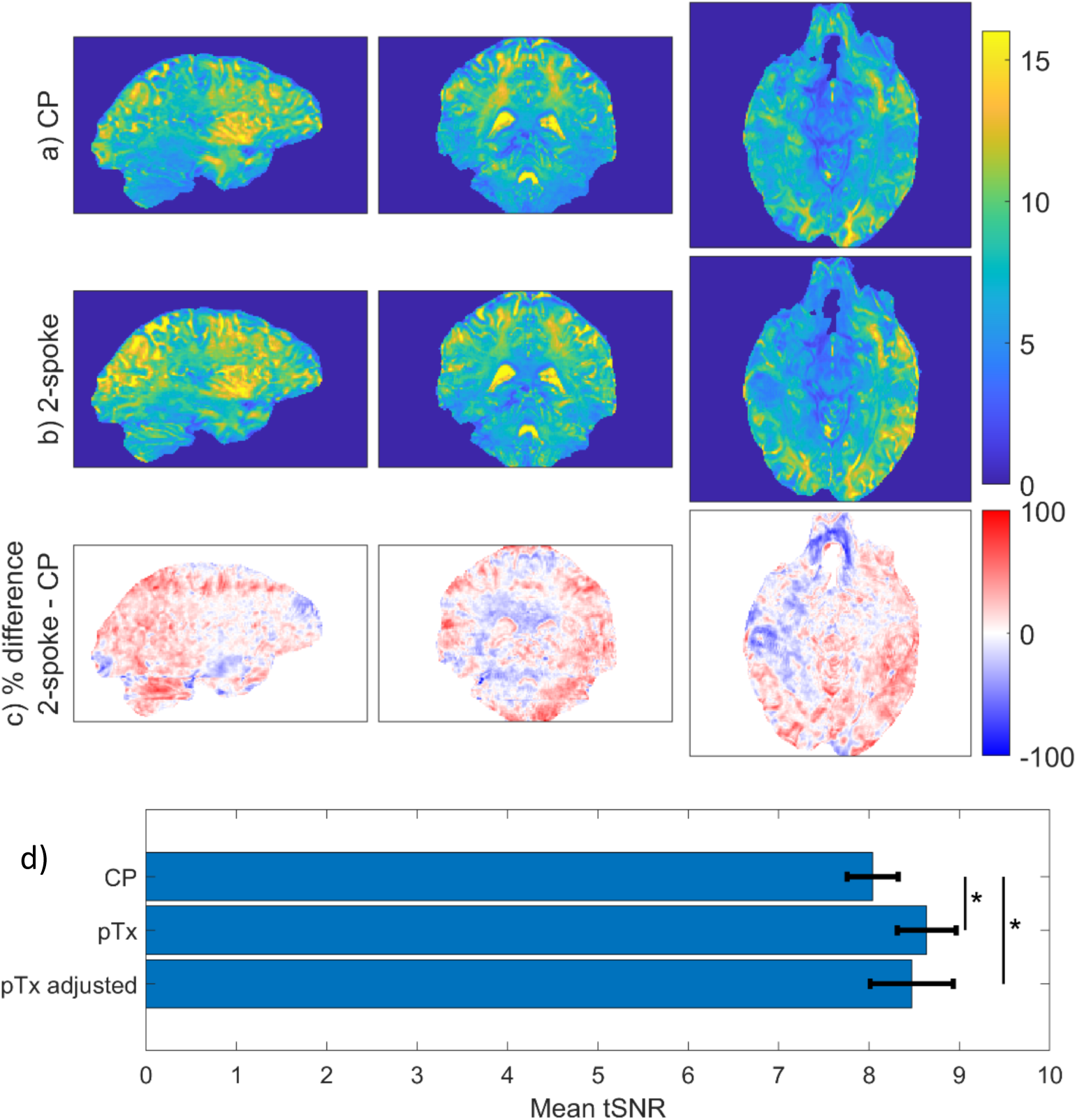
(a-c) Comparison of the temporal SNR between the CP and 2-spoke pTx scans. The tSNR is calculated from 10 repeated measurements of the no diffusion (b=0) images. (d) The whole brain mean tSNR across all subjects. ‘pTx adjusted’ divides the measured tSNR by sqrt(TR_pTx_/TR_CP_) to account for the lengthened TR in some subjects. pTx 2-spoke VERSE acquisitions show 5.4% increase in tSNR (P=0.004, T(5)) after adjustment.

As a result of the higher tSNR, the dynamic pTx pulses produced cleaner and more defined fractional anisotropy maps especially in the inferior parts of the brain as shown in Figure 4, where the CP data produced substantially more noise and the pTx images show fiber details more clearly.

**Figure 4.**
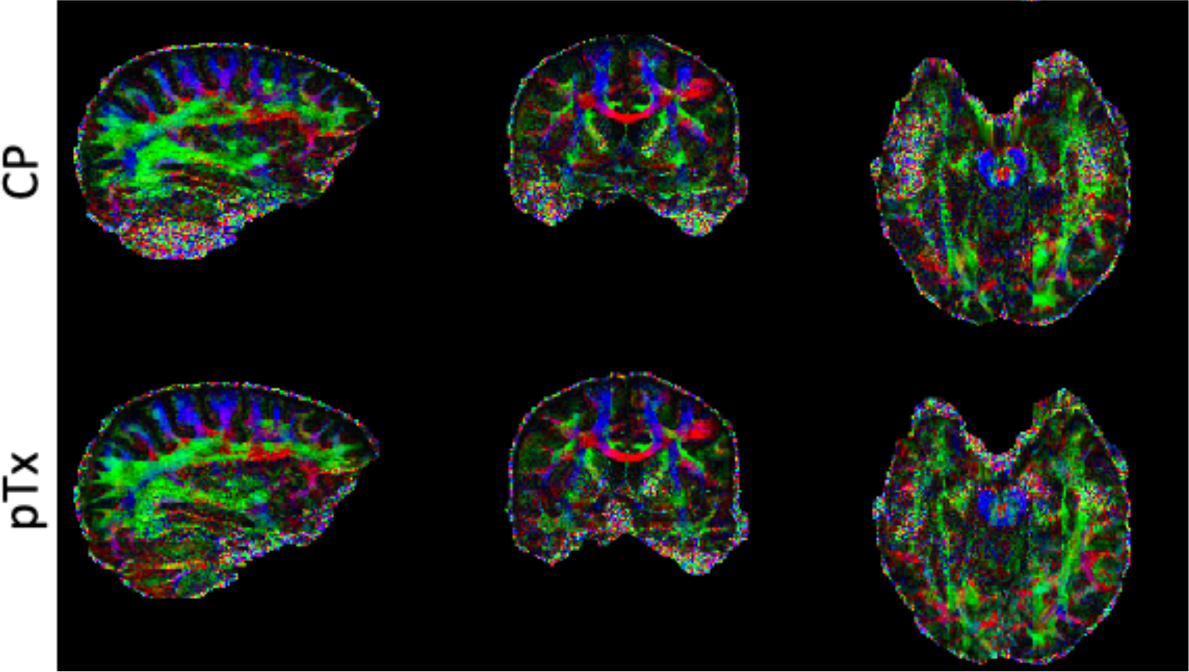
Fractional anisotropy color-coded by the principal diffusion direction (red - left-right; green – anterior-posterior; blue – superior-inferior) in one subject. The pTx 2-spoke data are less noisy and show better defined fiber orientations in the cerebellum, occipital lobe and inferior temporal lobes.

To quantify the improvement in diffusion tensor fitting, the first fiber orientation dispersion (uncertainty)[34] was obtained from ‘bedpost’ and transformed to the MNI space for all subjects. This is plotted in Figure 5. The pTx 2-spoke scans reduced the fitting uncertainty over the entire periphery of the brain, and especially in the cerebellum and inferior temporal lobes. The center of the brain showed some increase in the fitting uncertainty. This is again consistent with the differences in the flip angle patterns. Averaged the whole brain and all subjects, the principal fiber orientation uncertainty decreased by 6.2% (2-spoke VERSE 0.0655 vs CP 0.0703, *P*<0.001).

**Figure 5.**
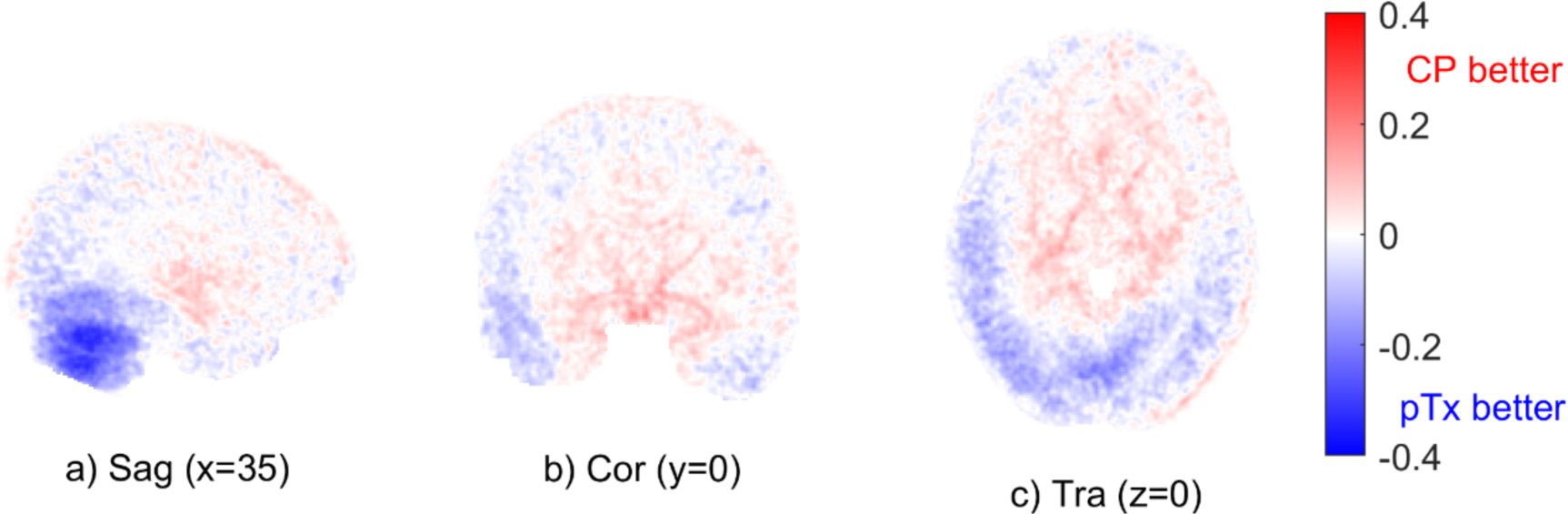
The difference in the first fiber orientation fitting uncertainties between CP and pTx acquisitions averaged across all subjects. Data of all subjects were transformed into MNI space for comparison. MNI coordinates are shown. Over the whole brain, pTx 2-spoke decreased the fitting uncertainty by 6.2% (P<0.001, T(5)).

This result is also reflected in the crossing fiber estimation performance. Figure 6 shows the second fiber volume fraction for all subjects, where the pTx data can increase the amount of resolved second fibers in the cerebellum, temporal lobes and all around the edge of the brain. The mean second fiber volume fraction increased by 19% from 2.31% to 2.74% (P=0.003, T(5)), with the second fiber orientation uncertainty showing a 1.3% decrease (2-spoke VERSE 0.139 *vs* CP 0.141, *P*=0.01).

**Figure 6.**
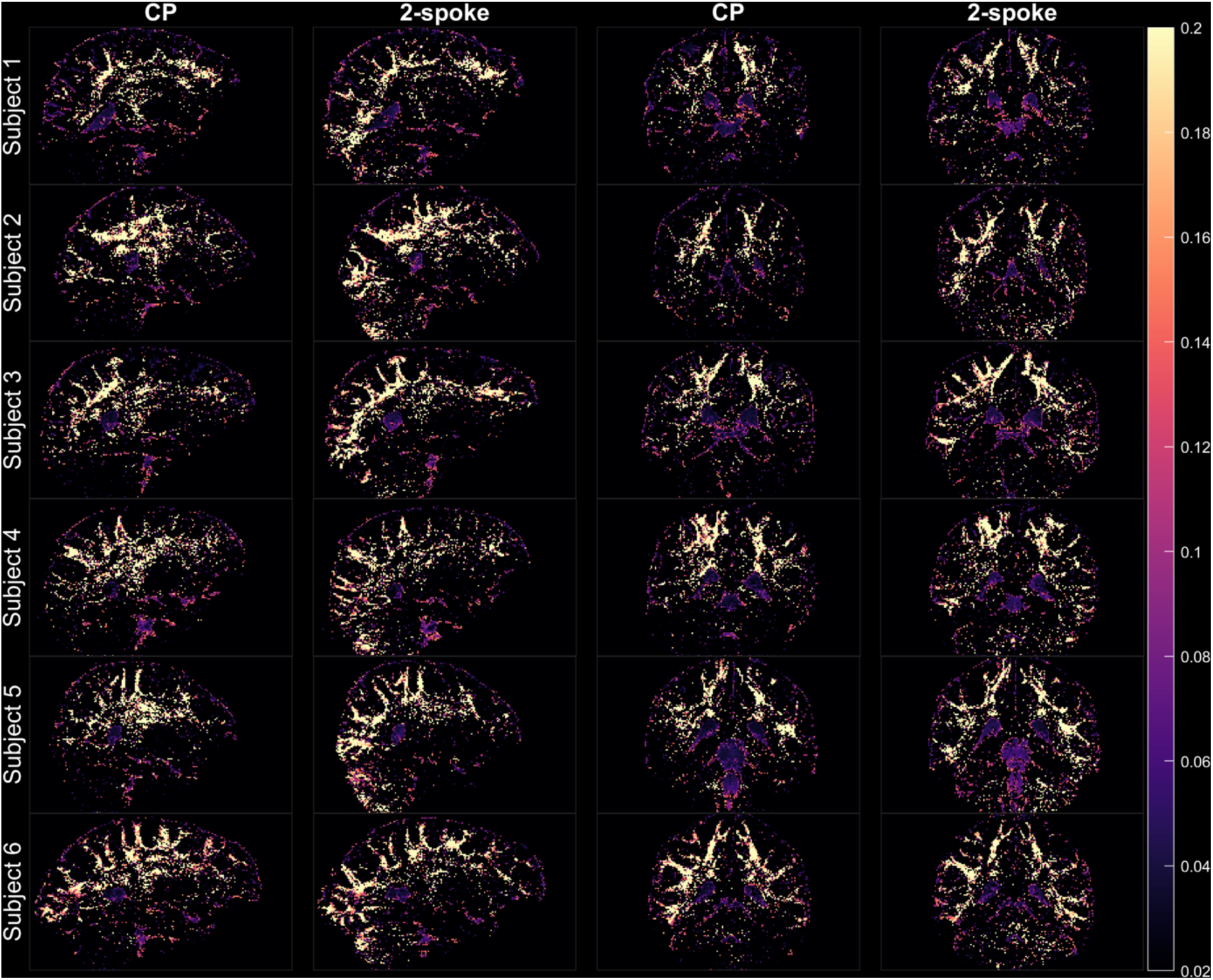
The second fiber fractional volume map plotted for all subjects. Color scale between [0.02 0.2]. pTx data allow more crossing fibers to be resolved around the periphery of the brain. The whole brain mean second fiber volume fraction increased by 19% from 2.31% to 2.74% (P=0.003, T(5)).

## Discussions

### Performance comparison

We have demonstrated the use of dynamic pTx pulses for dMRI acquisition in human brain. Our 2-spoke pTx VERSE pulses obtained more homogeneous spin-echo magnetization across the whole brain, improving the signal around the periphery of the brain, especially in the cerebellum and inferior temporal lobes. The data show that the changes in the flip angle patterns, the b=0 signal intensity, tSNR and the diffusion fitting uncertainties are consistent.

We chose to compare the performance of CP and 2-spoke dynamic pTx pulses by using the Nova 8Tx32Rx head coil throughout to eliminate any bias caused by the coil. We have shown elsewhere that the Nova 8Tx32Rx head coil achieves somewhat higher tSNR than the widely used Nova 1Tx32Rx head coil.[36] Therefore, we report a conservative estimate of the improvement for moving to pTx which in practice would also include an additional improvement by switching to the Nova 8Tx32Rx coil.

We used the similar acquisition parameters as the HCP 7T diffusion protocol to reflect the representative application of diffusion MRI at 7T. The number of diffusion directions and the total volumes acquired are reduced, so that the CP vs pTx comparison can be performed within a single session to minimize subject motion and avoid any other effects that might affect session-to-session reproducibility. For the same reason, we did not use dielectric pads for the CP scans to avoid repositioning the subject when removing the pads for the pTx scans.

The diffusion scans were performed with a pTx enabled sequence with minimum alteration from the vendor’s product sequence. As such, we did not implement some specific features used in the HCP diffusion sequence, such as the GRE GRAPPA autocalibration scans with the same ramp sampling as the EPI readout. This means that our data quality may not be as good as the original HCP protocols, but this does not affect our goal to compare performance for pTx vs CP pulses.

We chose to split the whole brain into 8 slabs for pulse optimization to better account for the B_0_ and B_1_^+^ variations especially in the inferior parts of the brain. We retrospectively tested the effect of number of slabs on predicted pulse RMSE (Supplementary Information Figure S1), showing that 8 slabs is a reasonable compromise albeit that for future studies it would be possible to reduce optimization time by merging slabs in the superior and middle parts of the brain.

### SAR and TR

Previous work suggested that designing pulses in the sagittal or coronal orientations can be more SAR efficient,[15] but we have attempted the pTx pulse design in the more usual transverse direction. Our results demonstrate that this is still a viable solution, but our pulse designs were limited by SAR.

In the pTx mode, with the Nova 8Tx32Rx head coil, the vendor enforces a 6-minute average power limit of 8 W total and 1 W per channel through a ‘diagonal’ virtual observation point (VOP) matrix in the IEC First Level mode. Besides, for extra safety, the current online VOP supervision system in the Terra pTx framework adds up the maximum channel power for each pulse, instead of adding up the channel power for all pulses before taking the maximum. This results in a further 10%-15% overestimation of SAR exposure.

In order to achieve a similar TR as the CP scan under the SAR constraints, we had to reduce the flip angles to 80° and 160°, and to choose a relatively high Tikhonov regularization factor λ=0.025 for the high flip angle optimizer, which limits the performance of the pulse. The VERSE conversion reduced the total SAR by 52% (inter-subject mean), at a cost of increased sensitivity to off-resonance.

With these limitations, we scanned at approximately 100% SAR level at the same TR as the CP mode acquisition with the Nova head coil. Specifically, for a typical set of pulses: the 110° fat saturation pulses account for approximately 30% SAR; the 80° excitation pulses, averaged over the 8 sub-volumes, account for ca. 19%; the remainder of the SAR allowance is taken by the 160° refocusing pulses, ca. 51%. The detailed pulse energies for each subject are presented in Table 1. Note that although the fat saturation pulses always had CP amplitudes and phases, they are subjected to the stricter pTx SAR limits when other pulses in the sequence utilize pTx. We were not able to use the gradient reversal method[37] or the low-SAR method[38] alone to suppress the fat signal, because the VERSE pulses disrupt the relationship between ΔB_0_ and slice selection gradient, and can generate significant fat artefacts.[39]

However, there is reason to believe that the current vendor-provided VOP limit may be overly conservative. NeuroSpin reported a SAR safety workflow which allows scanning with the Nova head coil with 16 W total, 3 W per channel limits and custom VOPs with no incidents on VB17 software version.[40, 41]

To investigate whether our sequence would perform better when SAR exposure is overestimated less severely, we ran equivalent scans on the Glasgow 7T Terra scanner which was equipped with a different 8Tx32Rx head coil for which full (non-diagonal) VOP models are available. This showed that all scans ran at <50% of the IEC first level SAR limit. The extra SAR allowance therefore opens the possibility to scan the same 2-spoke VERSE designs with at least multiband factor 2 at minimum TR. Alternatively, the extra SAR budget could be used to increase the flip angles to 90° and 180° and increase the magnetization quality, or it may be used to decrease the level of VERSE modification to improve resilience to off-resonance. Some examples of these potential improvements are predicted in Figure 7.

**Figure 7.**
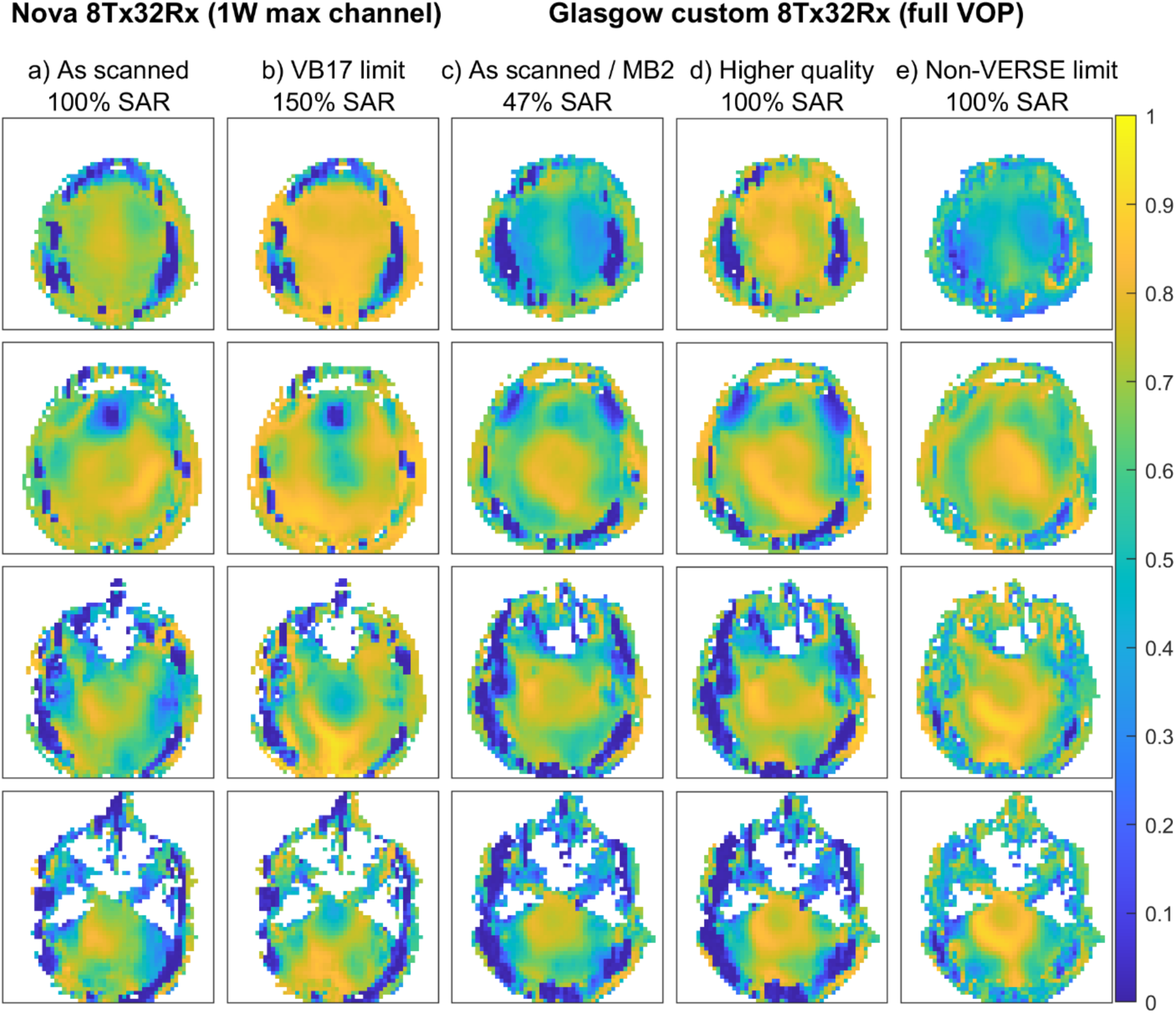
Predicted spin-echo intensity by Bloch simulation for several SAR limit levels for various cases a) 100% SAR level with 1W/channel limits (as provided with the Nova 8Tx32Rx coil on 7T Terra scanners with VE12 software). b) 100% SAR level with 1.5W/channel and 8W total (as provided with the Nova 8Tx32Rx coil on Magnetom 7T scanners with VB17 step 2.3 software). c) 50% SAR level, i.e. Normal Mode limit, for the custom coil that we used in Glasgow. This allows MB2 scans at minimum TR. d) 100% SAR level for the Glasgow custom coil. This shows how the extra SAR budget of the VOP model could be used to increase pulse design quality. e) Same, but without VERSE and with fat suppression disabled. This shows how a higher SAR budget could improve spin-echo uniformity by reducing the sensitivity to off-resonance. Note that these figures are plotted only to compare in broad terms the effect of the SAR budget and are not intended as a comment on the two coils.

### Bipolar spokes gradient delay

We chose to use monopolar spokes pulses in this study despite a longer excitation duration, because it has been shown[42] that bipolar spokes pulse performance are negatively impacted by imperfections in MR system gradient-RF timing. This phenomenon degrades signal in the inferior parts of the brain, due to an effective phase difference between the first and second spokes in the bipolar spokes pulse that is proportional to the slice distance from the isocenter.

We measured the RF—gradient timing difference on our system with two approaches: the double-navigator method proposed by Tse *et al.*[42] gave a delay of 2.5 μs; and the zero-flip-angle method of Gras *et al.*[43] gave a delay of 3.1 μs. Such delays can have a significant impact on flip angle homogeneity.

Although not specified in the original paper, we noticed that the excitation flip angle has an impact on the accuracy of the double-navigator method. As the flip angle increases, the experimentally measured timing delay is not consistent. To demonstrate this, we performed Bloch simulations of this double-navigator sequence with varying flip angles without any timing delays. A sinc pulse with TBW=4 and Hamming window was used. The slice selection gradient results in a 2 mm slice being excited, and the simulation grid had a resolution of 0.05 mm in the slice selection direction. The transverse magnetization was summed to simulate the signal seen by the receive coil. The sequence diagram and the simulated signals are shown in Figure 8a.

**Figure 8.**
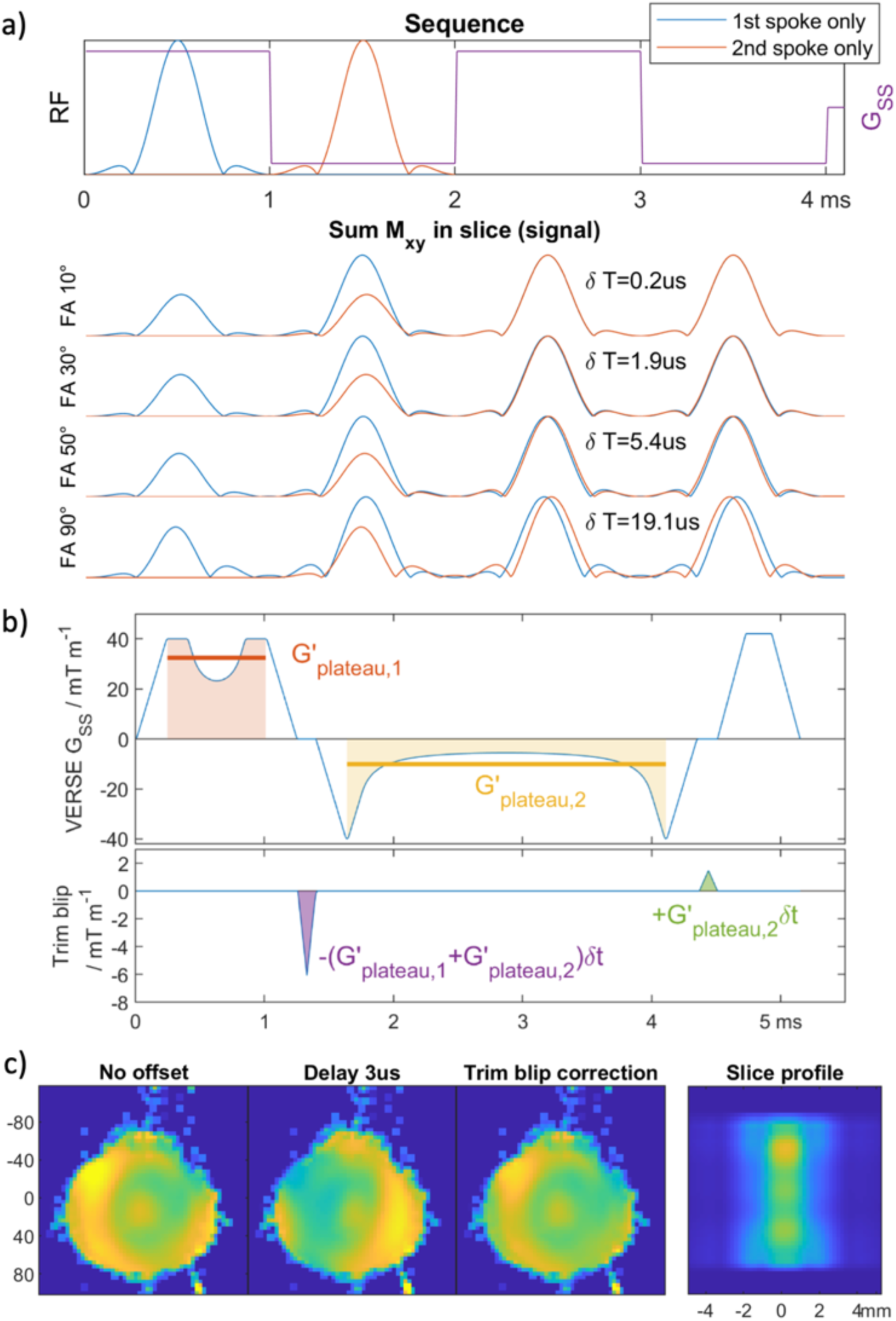
a) The double-navigator sequence (Tse et al.[42]) to determine the RF-gradient timing offset, and the Bloch simulated signal with varying flip angles. b) The calculation of the trim blip moment to correct for the timing delay in bipolar spokes as introduced in Jamil et al.[44] for VERSE pulses. The effective gradient strength is calculated with the slice selection moment divided by the corresponding ‘flat top’ time. c) Bloch simulation of the transverse magnetization of VERSE excitation pulses on a phantom, with 3 µs gradient delay at z = 8 cm from the isocenter. The magnetization pattern distortion caused by the timing delay is corrected by the trim blips, but the slice profile is smeared out and can not be mitigated with either monopolar spokes or the trim blip correction.

Even without an actual timing delay in the system, the measured echo peaks can be almost 40 µs apart (calculated timing delay 19.1 µs) just due to the asymmetric magnetization response. Therefore, it should be noted that the excitation flip angle needs to be small when using the Tse double-navigator method to minimize the systematic error.

We sought solution to these offsets using the method of gradient trim blips,[44] where small slice-selection gradient blips between the spokes are added to correct for the phase difference. We noted that the VERSE pulse does not have a flat gradient plateau, so the optimal blip moment should be found from the average gradient strength ((total slice-selection moment – ramp moment) / (pulse time – ramp time)). This is illustrated in Figure 8b.

To validate this formula, we conducted Bloch simulations of a phantom shown in Figure 8c, at z = 8 cm from isocenter with 3 µs gradient delay. The above trim blip moment is shown to restore the magnetization pattern of the 2-spoke VERSE pulses.

However, due to the nature of the VERSE pulses, the timing offset also causes an error in the slice-selection direction where the slice profile is smeared out (also shown in Figure 8c). This cannot be mitigated by using either monopolar pulses or the trim blip correction method. Our simulation results show an approximately 8% signal coming from outside the desired slice with the timing-corrected VERSE pulse compared to a system with no delay. This value should represent an upper bound for this effect for whole-brain imaging.

The full implementation of this gradient trim blip method for bipolar spokes VERSE pulses is beyond the scope of this work and will be addressed in a future study.

## Conclusions

We have successfully applied subject-specific 2-spoke dynamic pTx pulses for whole-brain diffusion MRI. Compared to matched scans with circularly polarized (CP) pulses, whole-brain flip angle RMSE reduced and whole-brain tSNR increased. Diffusion tensor fitting of the pTx data had lower uncertainty.

## CRediT authorship contribution statement

Minghao Zhang: Software, Methodology, Investigation, Formal analysis, Visualization, Writing - Original Draft. Belinda Ding: Software, Methodology, Writing - Review & Editing. Iulius Dragonu:

Conceptualiztion, Software, Resources. Patrick Liebig: Software, Resources. Christopher Rodgers: Conceptualization, Methodology, Writing - Review & Editing, Supervision, Funding acquisition.

## Acknowledgements

The authors thank Prof David Porter, Dr Sydney Williams, Dr Shajan Gunamony, Dr Graeme Keith, Steven Winata and Janhavi Ghosalkar for facilitating the scans in Glasgow, and Dr Robin Heidemann for helpful discussions.

## Supplementary information

**Supplementary Information Figure S1.**
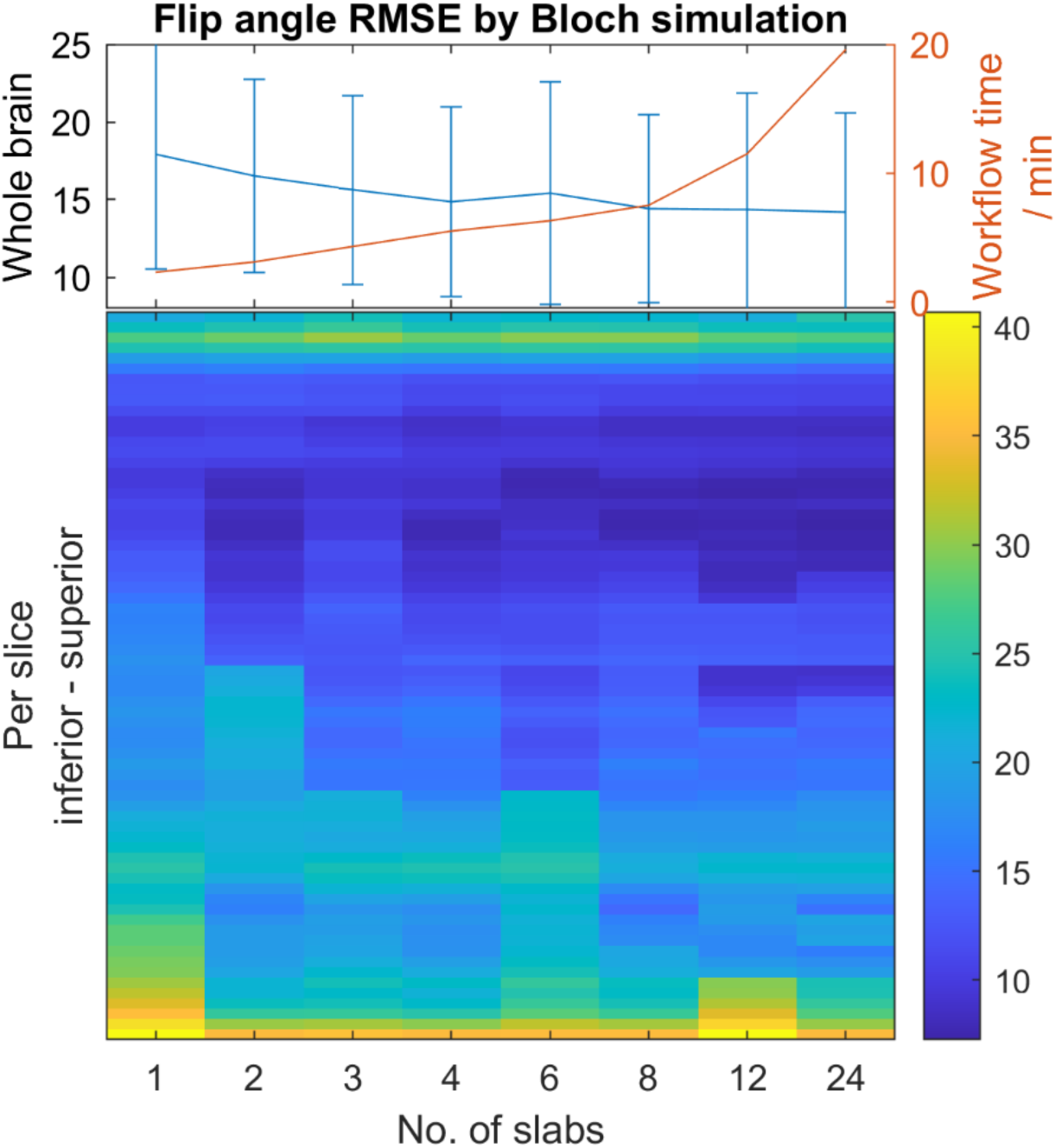
Flip angle root mean squared error (in °) for the 80° excitation pulse computed from Bloch simulation results in one subject, (a) over the whole brain and (b) per transverse slice. The workflow time consists of the excitation and refocusing pulse design, visualization and data transfer to and from the scanner. Our choice of 8 slabs is a good compromise between flip angle accuracy and optimization time.

