## Supplementary Information for "Dynamic parallel transmit diffusion MRI at 7T"

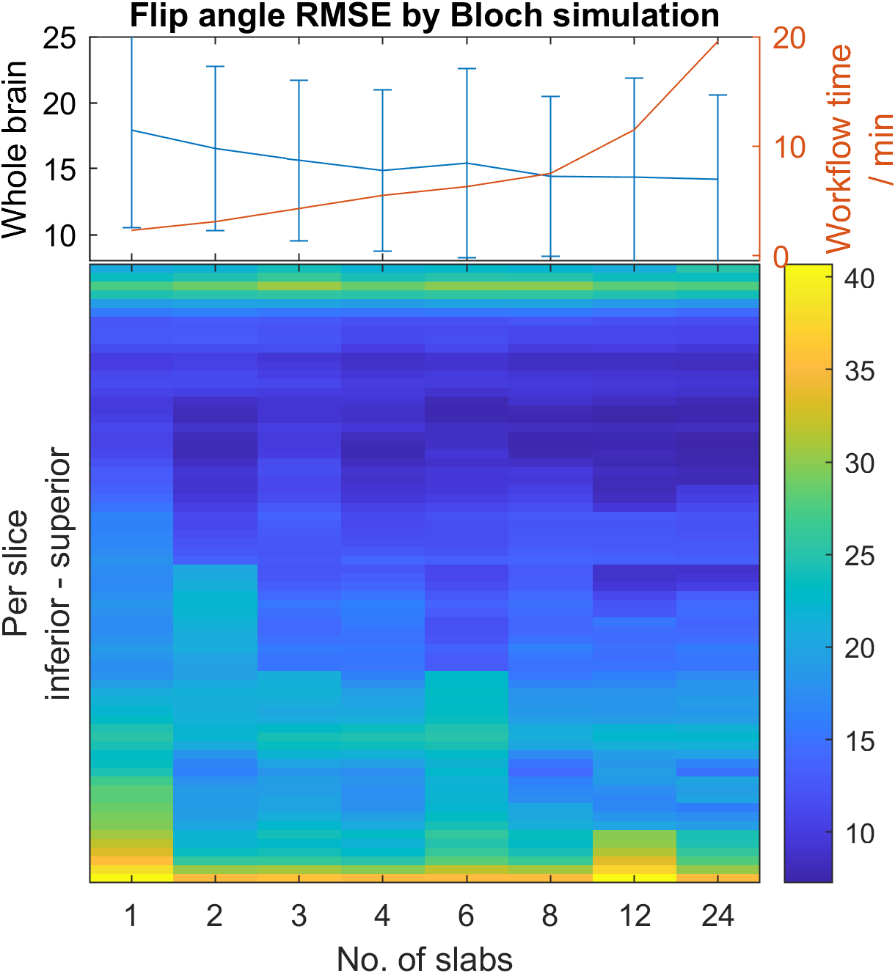

Supplementary Information Figure S1 Flip angle root mean squared error (in °) for the 80° excitation pulse computed from Bloch simulation results in one subject, (a) over the whole brain and (b) per transverse slice. The workflow time consists of the excitation and refocusing pulse design, visualization and data transfer to and from the scanner. Our choice of 8 slabs is a good compromise between flip angle accuracy and optimization time.
